# Sparse delivery of Synaptobrevin-2 to neurons using extracellular vesicles

**DOI:** 10.1101/2025.11.04.686638

**Authors:** Victoria Hannett, A. Alejandro Vilcaes, Siewert Hugelier, Qing Tang, Melike Lakadamyali, Natali L. Chanaday

## Abstract

Synaptobrevin-2 (Syb2) is an essential SNARE protein for neurotransmitter release and communication in the nervous system. We previously showed that Syb2 is also exchanged among neurons via extracellular vesicles (EVs). Host neurons can rapidly incorporate exogenous Syb2 into their synaptic vesicle cycle to support neurotransmitter release, however the endocytic mechanism of Syb2-containing EVs is unknown. Here, we use a fusion of Syb2 with the pH-sensitive GFP (Syb2-pHluorin) to track the incorporation of Syb2-containing EVs into neurons and the trafficking of exogenous Syb2 to synaptic vesicles at synapses. We determined that Syb2-containing EVs are endocytosed via Clathrin- and Dynamin-independent pathways. Moreover, Syb2-containing EVs are directly uptaken by axons and are rapidly incorporated into functional synaptic vesicles. These Syb2-pHluorin positive synaptic vesicles are endocytosed with either ultrafast (<1s) or fast (∼1-3s) kinetics during synaptic transmission, suggesting limited diffusion and high fidelity in the fast retrieval of synaptic vesicle molecules immediately after fusion. This work introduces a novel application of EVs as vehicles to deliver fluorescent molecules in a neuron-specific, targeted manner to investigate protein transport mechanisms without overexpression artifacts. Our findings expand our understanding of the mechanisms EVs use to enter neurons.

## INTRODUCTION

Neuronal communication spans a wide range of timescales and distances. Fast information processing in neuronal circuits is performed via electrical impulses and chemical transmission at synapses, while slower trophic and homeostatic signals are transferred via secretion of neuromodulator molecules and, the latest added players, extracellular vesicles (Blanchette & Rodal, 2020; Nieves Torres & Lee, 2023; Sullivan et al., 2025). A common element regulating all these fast and slow communication modes is the vesicular protein Synaptobrevin-2 (Syb2 or vesicle-associated membrane protein 2 – VAMP2). However, how Syb2 is transferred between extracellular and intracellular compartments is unknown.

Syb2 was first discovered in synaptic vesicles from rat brain where it is essential for the fusion of synaptic vesicles and the subsequent release of neurotransmitters (Link et al., 1992). It was later discovered that Syb2 is a SNARE (soluble NSF attachment receptor) protein that, together with its target SNARE partners SNAP-25 and Syntaxin-1, catalyzes the fusion of vesicles with the target membrane (Hayashi et al., 1994; Pevsner et al., 1994). In this manner, Syb2 mediates the release of secretory vesicles and the surface delivery of plasma membrane proteins in multiple cell types, including neuronal, glial, endocrine, adipose and immune cells (Actis Dato et al., 2018; Bakr et al., 2021; Chang et al., 2015; Chen et al., 2023; Lam et al., 2022; Martin et al., 1998; Regazzi et al., 1995; Schoch et al., 2001; Uriarte et al., 2011). Syb2 has been proposed to be retrieved from the target membrane via multiple endocytic pathways, including Clathrin-dependent and independent endocytosis (Cousin, 2021; Gordon et al., 2011; Koo et al., 2015; Koo et al., 2011; Renard et al., 2015). Moreover, Syb2 couples exocytosis to endocytosis of synaptic vesicles during synaptic transmission, ensuring fast recycling of vesicles and maintaining the efficacy of neurotransmission (Chanaday & Kavalali, 2021; Deák et al., 2004). Recently, Syb2 was found to be itself secreted to the extracellular space of neurons via extracellular vesicles (EVs), which could be a new non-synaptic mechanism of neuron communication (Vilcaes et al., 2021).

Thus, Syb2 is involved in multiple forms of trafficking and secretion. However, majority of previous research on Syb2 trafficking was performed by overexpressing fluorescently tagged versions of Syb2 in neurons, which causes abnormal subcellular distribution and recycling of Syb2, including higher plasma membrane levels and impaired endocytosis (Gordon et al., 2016; Pennuto et al., 2003). Here we take advantage of our previous discovery showing Syb2 secretion via EVs and use these EVs to deliver tagged Syb2 to neurons. Using this novel approach, we investigate the endocytic mechanism of EVs carrying Syb2 and the transport of this exogenous Syb2 after incorporation into neurons. We also demonstrate an application of this approach to study the trafficking of Syb2 at synapses in a more “native” environment. The advantages of this new delivery method are absence of overexpression artifacts, low background fluorescence and low toxicity – since transfection is not needed.

## METHODS

### Dissociated hippocampal cultures and isolation of extracellular vesicles

Dissociated hippocampal cultures from embryonic day 17-19 mice (SNAP25 KO) (Bronk et al., 2007) or postnatal day 0-2 Sprague-Dawley rats of both sexes were used for all the experiments. All experiments were performed following protocols approved by the University of Pennsylvania Institutional Animal Care and Use Committee.

Bilateral hippocampi were digested using 10 mg/mL trypsin and 0.5 mg/mL DNAse for 10 min at 37°C. Tissue was carefully dissociated using a P1000 pipette and cells were plated onto Matrigel (Corning, catalog # 354234) coated plastic bottom plates (for EVs isolation) or 8-well LabTek (for microscopy). Basic growth medium consisted of MEM medium (no phenol red), 5 g/l D-glucose, 0.2 g/l NaHCO3, 100 mg/l transferrin, 5% of fetal bovine serum, 0.5 mM L-glutamine, 2% B-27 supplement, and 2–4 μM cytosine arabinoside. Cultures were kept in humidified incubators at 37°C and gassed with 95% air and 5% CO_2_.

For EVs isolation, neuron cultures at DIV 12-14 were used. 50% of culture media was replaced and 30% of fresh media was added every 24h. After 60 h, culture supernatant was collected and cleared by serial centrifugations as previously described (Vilcaes et al., 2021). The resulting supernatant was used for EVs isolation by size exclusion chromatography using qEV1/35mm columns from IZON (Catalog # IC1-35) and following the procedure suggested by the manufacturer. Nanoparticle Tracking Analysis A NS300 NTA machine (Malvern NanoSight NS300) was used to analyze isolated EVs.

For all the experiments, a concentration of 7.5×10^8^ particles/ml of EVs was used, this concentration is equivalent to the normal concentration of EVs in the supernatant of dissociated neurons (Vilcaes et al., 2021).

### Cloning and lentiviral infection

Lentivirus for Syb2-pHluorin expression was produced in HEK293T cells (ATCC, catalog # CRL-1573) by cotransfection of pFUGW-Synaptobrevin2-pHluorin vector (Ramirez et al., 2012) and 3 packaging plasmids (pCMV-VSV-G, pMDLg/pRRE, pRSV-Rev - AddGene # 8454, 12253 and 12251) (Dull et al., 1998; Stewart et al., 2003) using FuGENE 6 transfection reagent (Promega, catalog # E2692). Fresh, cleared supernatants containing lentiviruses were used for infection of days in vitro (DIV) 4 hippocampal neurons.

### Incubation with EVs, endocytic markers and Dyngo-4a

Neuron culture media was replaced with 120 µL of Neurobasal solution containing 7.5×10^8^ particles/ml of Syb2-pHluorin-containing EVs, transferrin-Alexa647 (TF; Invitrogen, catalog # T23366), BODIPY-lactosylceramide (LacCer; Invitrogen, catalog # B34402) and/or 80 µM 3-hydroxy-2-naphthalenecarboxylic acid (Dyngo-4a; Abcam, catalog # ab120689), and incubated for 10, 30 or 60 min at 37°C in humidified incubators with 5% CO_2_. For the Dyngo-4a negative control, a similar volume (5 µl) of DMSO vehicle was added to the aforementioned mixture.

### Immunofluorescence

Neuron cultures were fixed with 1% PFA and 4% sucrose in DPBS and permeabilized using 0.01% digitonin in DPBS. After blocking with 3% BSA, primary antibodies against GFP (1:100) (Invitrogen, catalog # A11122), Syn1 (1:500) (Synaptic Systems, catalog # 106011) and/or MAP2 (1:1000) (Synaptic Systems, catalog # 188004) were diluted in blocking buffer and incubated overnight at 4 °C in a humid chamber. Secondary antibodies were incubated for 1h at room temperature. For STORM, secondary antibodies were conjugated with the activator– reporter dye pairs (anti-mouse conjugated to Cy3b and Alexa 405 at 1:100, anti-rabbit conjugated to Alexa 647 and Alexa 405 at 1:100) by the laboratory of Dr. Melike Lakadamyali following previously reported procedures (Bates et al., 2007).

### STORM experiments

Three controls were performed in independent experiments presented in Figures 2 and 3: A negative control consisting of dissociated neurons stained only with secondary antibodies (to control for secondary antibody specificity). A second negative control of cultures not treated with EVs and labeled with anti-GFP antibody (to control for primary antibody specificity). And a positive control using neurons that overexpress Syb2-pHluorin via lentiviral infection (necessary control to tune colocalization analysis).

**Figure 1.**
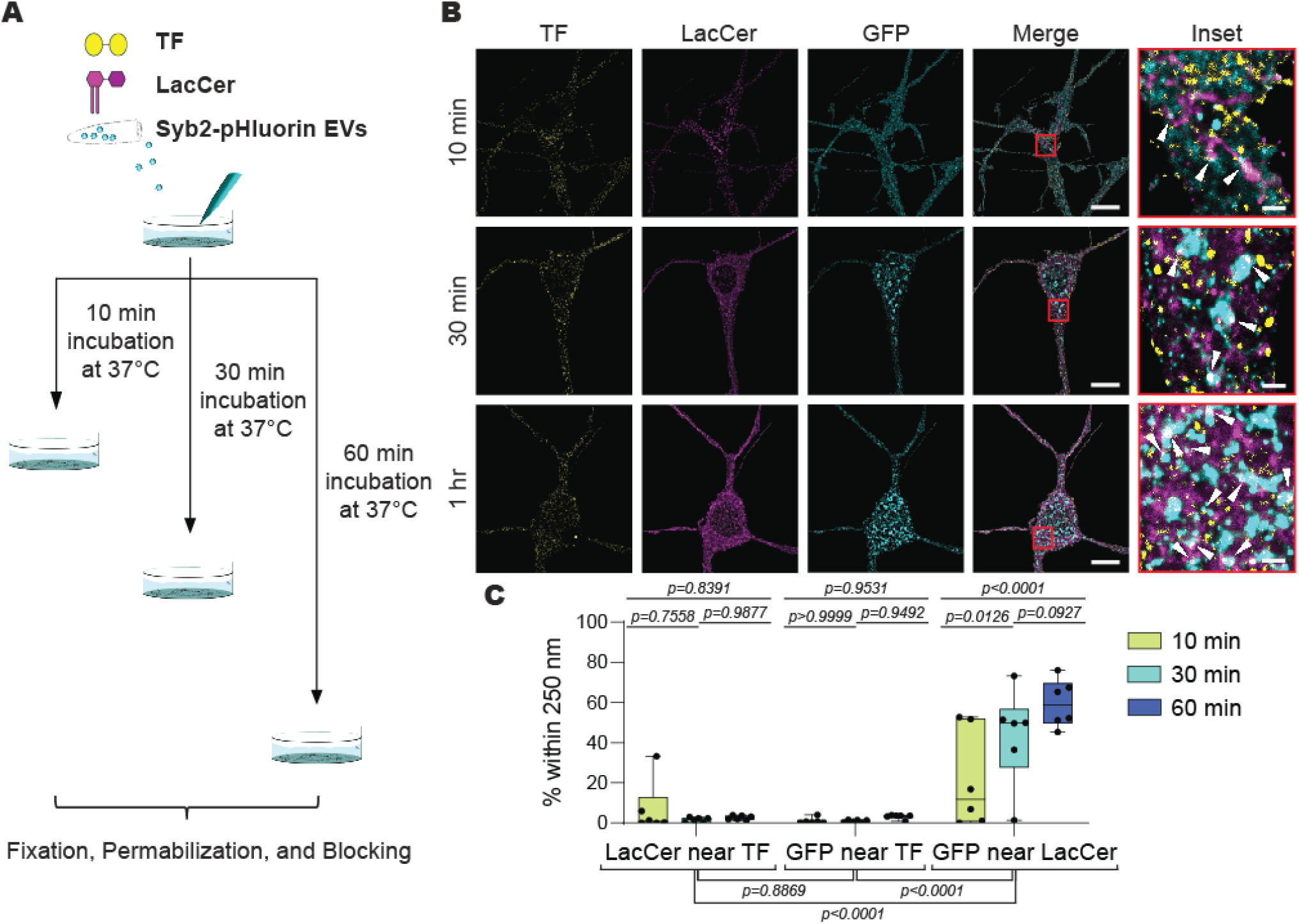
EV-delivered Syb2 trafficking converges with LacCer. **A**. Experimental design. **B**. Representative confocal images of neuronal cell bodies immunofluorescently labeled with TF (yellow), LacCer (magenta) and anti-GFP (cyan). Z-stacks of images are shown, presented as the sum-of-slices projection of the volumetric data. White scale bars = 10 µm. Inset white scale bars = 1 µm **C**. Percentage of fluorescent objects within 250 nm across two fluorescent labels, following all incubation times. To compare group-wise differences, two-way ANOVA: incubation time factor, *F=3*.*934, p=0*.*0266*; cluster proximity factor, *F=55*.*00, p<0*.*0001*. Each dot represents the average of all neuron somas from one image.

**Figure 2.**
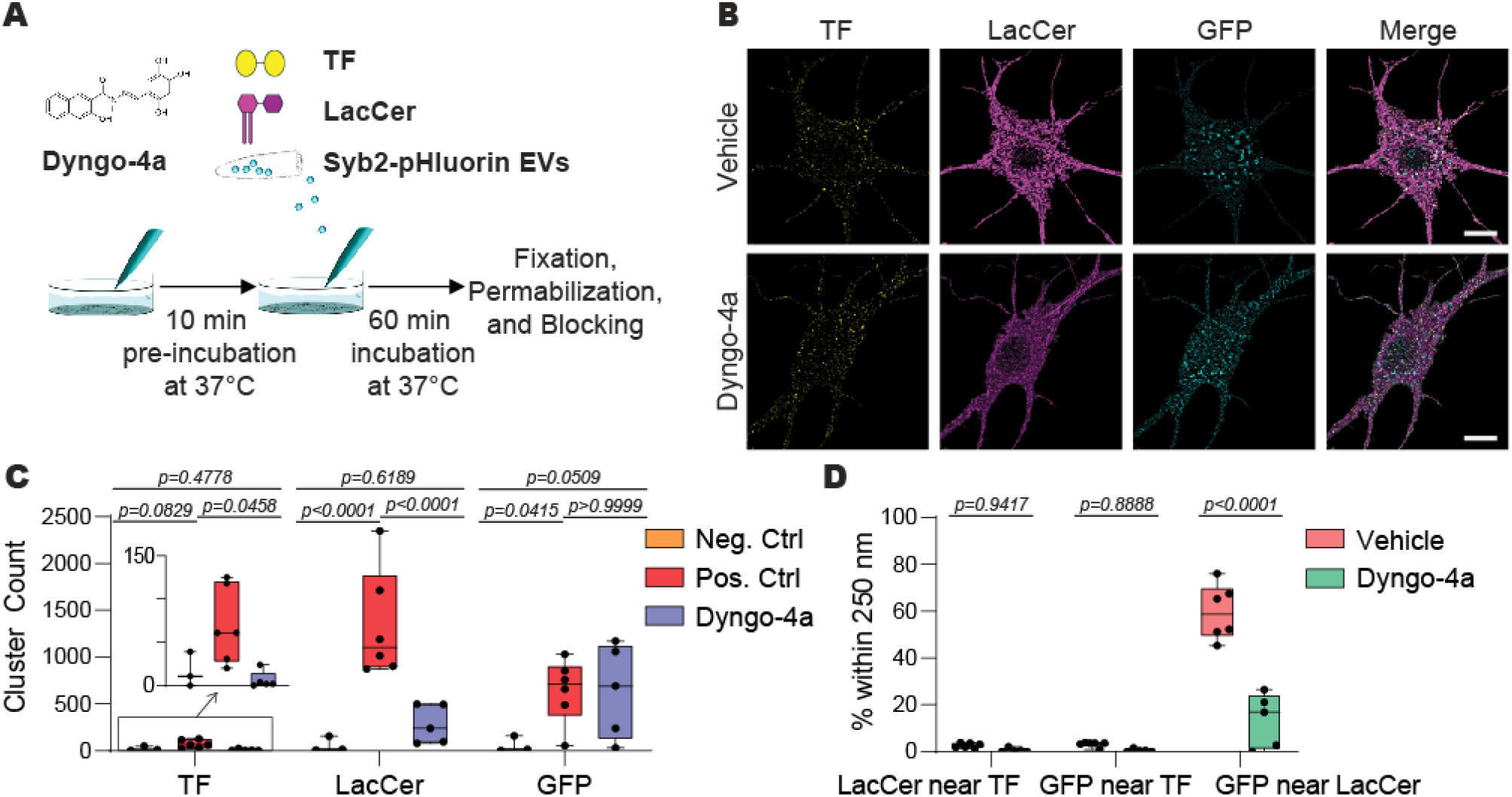
Syb2-containing neuronal EVs are endocytosed independently of Clathrin and Dynamin. **A**. Experimental design. **B**. Representative confocal images of neuronal cell bodies immunofluorescently labeled with TF (yellow), LacCer (magenta) and anti-GFP (cyan). Z-stacks of images are shown, presented as the sum-of-slices projection of the volumetric data. Scale bars = 10 microns. **C**. Number of TF, LacCer, and Syb2-pHluorin EV clusters in negative control (NC), positive control (PC), and Dyngo-4a-treated groups. Two-way ANOVA: experimental group factor, *F=12*.*76, p<0*.*0001*; cluster identity factor, *F=9*.*388, p=0*.*0006*. **D**. Percentage of fluorescent objects within 250 nm across two fluorescent labels, following treatment with vehicle (DMSO) or Dyngo-4a. Two-way ANOVA: experimental group factor, *F=51*.*23, p<0*.*0001*; cluster proximity factor, *F=95*.*44, p<0*.*0001*. Each dot represents the average of all neuron somas analyzed from one image.

**Figure 3.**
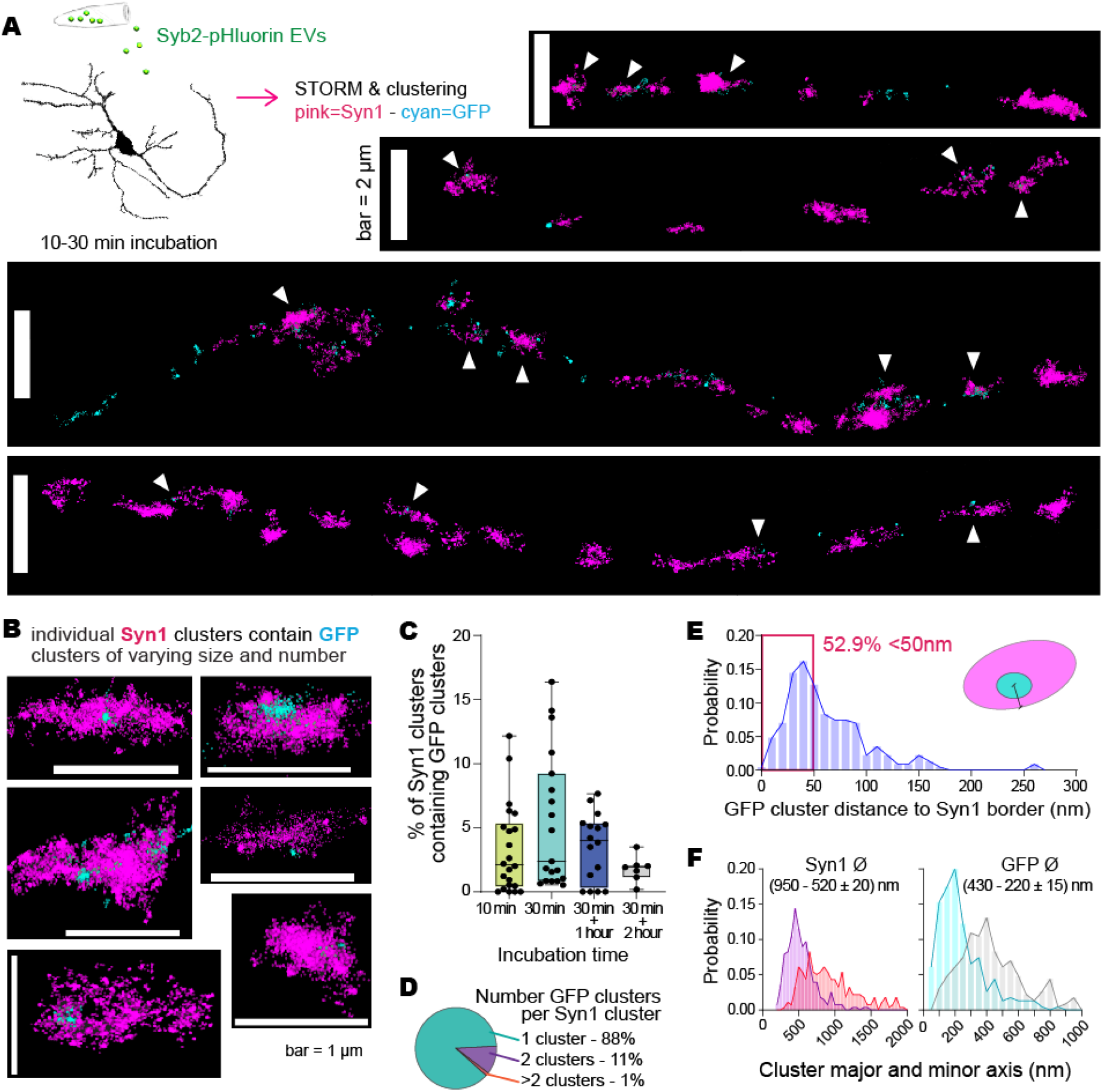
Syb2-containing neuronal EVs are uptaken by axons. **A**. Experimental design and representative STORM images showing Syn1 clusters in en passant axons (pink) and GFP clusters (corresponding to Syb2-pHluorin, cyan). **B**. Individual Syn1 presynaptic boutons containing GFP signal. **C**. Percentage of Syn1 boutons containing GFP signal. **D**. Number of GFP clusters per Syn1 cluster. **E**. Distance of GFP clusters to border of Syn1 cluster. **F**. Diameters of Syn1 (left) and GFP (right) clusters. Each dot represents one image, data from 3 independent cultures.

STORM images were acquired with a STORM microscope from ONI (Nanoimager S) using a 100X, 1.4 NA oil-immersion objective and a sCMOS Hamamatsu Orca Flash camera. To promote photoswitching of the fluorophores, a buffer containing 10 mM cysteamine MEA, 0.5 mg/ml glucose oxidase, 40 mg/ml catalase, and 10% glucose in PBS was used. The 640 and 561 lasers were used at 50% power, and the 405 laser was used at 5-8% or 15% power, respectively. 30,000 images were acquired with 10 ms exposure time for each channel (647 channel was always acquired first). STORM image localizations were obtained using Nanoimager software (ONI) and analyzed using custom-written MATLAB codes. In brief, clustering of the localizations was performed using Voronoi segmentation with the same threshold for all experiments (threshold was set using the negative controls), and clusters with less than 5 localizations were filtered out (there were no clusters detected in the negative controls). The Syn1 clusters were used as ‘reference’ data for the colocalization analysis, and the GFP clusters as the ‘colocalization’ data (see https://github.com/melikelakadamyali/StormAnalysisSoftware). The code transforms the reference cluster into high definition ‘alphaShape’ objects and then considers a GFP cluster to be colocalized when there is at least 40% overlap between the GFP localizations and the reference cluster alphaShape object. Using these parameters, there was more than 85% overlap and colocalization between Syn1 clusters and GFP clusters in the positive control. For the GFP clusters that are colocalized, the distance metric reported in Figure 2E was calculated by determining the distance between the center of the colocalization cluster and the closest border point of the reference cluster.

Fiducial markers (1:10000 dilution) were added to each imaged sample and used to correct for drifting. Alignment of the two (Syn1 and GFP) channels was checked before performing colocalization analysis.

### Confocal microscopy and analysis

Confocal images were acquired using a Zeiss LSM 980 Confocal Microscope (Carl Zeiss) with a 63X (NA1.4) objective and a pixel size of 0.044 x 0.044 x 0.21 microns (x,y,z). Around 6-8 Z-stacks were acquired in different regions of each coverslip/sample. Deconvolution was performed using the Fiji/ImageJ plugin DeconvolutionLab2 (Sage et al., 2017). The Richards & Wolf 3D Optical Model was used to calculate the Point Spread Function (PSF) using appropriate acquisition parameters. Regions of interest (ROIs) were drawn around individual neurons in each image using the MAP2 channel and each ROI was deconvolved using the Richardson-Lucy algorithm. Then the Fiji/ImageJ plugin Distance Analysis (DiAna) (Gilles et al., 2017) was used to determine nearest neighbor distances between GFP (EV), LacCer, and TF signals. Thresholds for DiAna Labelisation were determined using a negative control (neurons incubated with secondary antibodies but without primary). The data was filtered to determine the number of positive objects that were within 250 nm distance compared to the total number of positive objects, this distance was defined as colocalization since it is the limit of resolution of our images.

### Live fluorescence imaging

Cultured hippocampal neurons were imaged in modified Tyrode’s buffer containing 6-cyano-7-nitroquinoxaline-2,3-dione (CNQX, 10 μM) and aminophosphonopentanoic acid (AP-5, 50 μM) to avoid recurrent network activity. Fluorescence was recorded using a Nikon Eclipse TE2000-U microscope (Nikon) and an Andor iXon+ back-illuminated EMCCD camera (Model no. DU-897E-CSO-#BV). For illumination, we used a Lambda-DG4 illumination system (Sutter Instruments) with a FITC emission filter. Images were acquired at 5 Hz. Circular regions of interest (ROI) of 2 µm diameter were drawn around local fluorescence maxima (putative presynaptic boutons) and measured using Fiji (NIH). Fluorescence peaks synchronous respect to the stimulation were detected and analyzed using MATLAB (Chanaday & Kavalali, 2018, 2021).

### Statistical Analysis

For all figures, the full distribution of all experimental data is shown. Statistical analysis was performed using GraphPad Prism software. For Figures 1 and 2, data was analyzed using 2-way ANOVA with Tukey multiple comparisons tests, and statistical details are provided in the figures and their legends. For Figure 5, histograms were fitted in MATLAB using a Gaussian mixture model with one to twenty components, and the model with the smallest Bayes information criterion (BIC) value was automatically selected as the best fit (Chanaday & Kavalali, 2021). All data results from 2-3 independent cultures. Efforts were made to minimize the number of animals used for the experiments.

## RESULTS

### Uptake of Syb2-containing EVs into hippocampal neurons occurs independently of Clathrin and Dynamin

We previously showed that dissociated hippocampal neurons secrete Syb2 via EVs, through a mechanism that depends on the tretaspanin CD81 (Vilcaes et al., 2021). Using EVs isolated from neurons expressing Syb2 tagged with pH-sensitive GFP – pHluorin – we found that neuronal EVs containing Syb2-pHluorin are preferentially internalized into neurons (Vilcaes et al., 2021), although the uptake mechanism is unknown. The preferred uptake pathways for EVs in multiple cell types seems to be endocytosis (Mulcahy et al., 2014), and a previous report suggested that EVs incorporation into cortical neurons occurs via Clathrin and Dynamin dependent endocytosis, specifically (Solana-Balaguer et al., 2023). Neurons and neuronal EVs are rich in glycosphingolipids, which are thought to be crucial for EVs secretion and EVs effects on target neurons (Monyror et al., 2025; Yuyama et al., 2014). Glycosphingolipids are retrieved from the surface mainly via Caveolin-mediated endocytosis (Nusshold et al., 2013; Singh et al., 2003), so it is possible that neuronal EVs also use this pathway for their uptake. Therefore, to determine the internalization route of Syb2-containing EVs into hippocampal neurons we quantified the colocalization of GFP – which labels the Syb2-pHluorin molecules delivered via EVs – with Transferrin-Alexa647 (TF), a classical marker of Clathrin-mediated endocytosis, and BODIPY-lactosylceramide (LacCer), a marker of Caveolin-dependent endocytosis. We incubated neurons with Syb2-pHluorin EVs, TF and LacCer for 10, 30 or 60 min at 37°C and then performed confocal microscopy (Fig.1A-B). Microtubule-associated protein 2 (MAP2) was used to visualize the neurons and individual cell bodies (or soma) were selected for analysis. Images were deconvolved and analyzed using the Fiji plugin DiAna (Gilles et al., 2017) to determine the nearest-neighbor distance between the different endocytic intermediaries. Objects closer than 250 nm were defined as colocalizing. We did not observe colocalization between TF and LacCer at any of the measured time points (only 2±1% overlap; Fig.1B-C), in agreement with a recent report proposing that Clathrin and Caveolin pathways may not converge to the same early endosomes in neurons (Shikanai et al., 2023). GFP-positive objects were also rarely seen near TF-positive endosomes, and instead were found close to LacCer (Fig.1B-C). Moreover, the proximity between GFP and LacCer objects increased as a function of time (Fig.1B-C), suggesting that Syb2-pHluorin delivered via EVs is trafficked through a pathway that converges to the same endosomal organelles as Caveolin cargoes.

Our data suggests that Syb2-containing EVs are internalized via a Clathrin-independent endocytic route in hippocampal neurons. Moreover, Syb2 from the EVs converges to the same endosomal intermediary as the Caveolin cargo LacCer, but their low colocalization at short time points suggests different uptake mechanisms. Endocytic pathways can be globally classified depending on their reliance on the membrane scission molecule Dynamin (Gundu et al., 2022; Rennick et al., 2021). Since both Clathrin and Caveolin pathways are Dynamin-dependent, we next speculated that the internalization of Syb2-containing EVs may be Dynamin-independent. To test this hypothesis, we used the specific inhibitor Dyngo-4a to block Dynamin-dependent endocytosis and incubated neurons with Syb2-pHluorin EVs, TF and LacCer for 60 min at 37°C (Fig.2A). While pretreatment of hippocampal neurons with Dyngo-4a abolished the endocytosis of TF and LacCer, it did not significantly affect the internalization of EVs-delivered Syb2-pHluorin (detected with anti-GFP antibody; Fig.2B-C). In consequence, the proximity between GFP and LacCer objects significantly decreased in the Dyngo-4a treated neurons (Fig.2D). Taken together, these results demonstrate that Syb2-containing EVs are largely incorporated into hippocampal neurons through a Clathrin- and Dynamin-independent pathway.

### Syb2-containing EVs are directly internalized into axons

We previously reported that Syb2-containing EVs increase spontaneous inhibitory neurotransmission in the target neuron as soon as 20-30 min after addition (Vilcaes et al., 2021). The data presented here also shows that after internalization into the neuron soma, Syb2 from the EVs is trafficked through an endosomal pathway that converges with Caveolin cargoes 30-60 min after internalization. That timescale of intracellular transport is incompatible with the fast functional effects in synaptic transmission we previously uncovered, suggesting that Syb2-containing EVs may be directly endocytosed at the synapse. Due to the narrow caliber of axons (<1 µm wide) we turned to super-resolution microscopy to determine the localization of EVs-delivered Syb2-pHluorin along axons of dissociated hippocampal neurons. Neurons were incubated with EVs for 30 min at 37°C, then fixed, immunostained and imaged via Stochastic Optical Reconstruction Microscopy (STORM; Fig.3). As previously, exogenous Syb2-pHluorin, incorporated into neurons from the EVs, was identified with anti-GFP antibody. Image segmentation was performed using a previously validated Voronoi tessellation method (Gyparaki et al., 2021) (clustering and colocalization parameters were determined using negative and positive controls, see Methods). In EV-treated neurons, small GFP-positive clusters were observed in structures resembling *en passant* axons (Fig.3A), usually colocalizing with the presynaptic marker Synapsin-1 (Syn-1; Fig.3A-B). Only a small fraction of presynaptic terminals contained GFP (<10-15%; Fig.3C), agreeing with our previous measurements using confocal microscopy (Vilcaes et al., 2021). Since we are limited to detection of GFP clusters rather than single GFP molecules we are likely underestimating the real number of presynaptic terminals containing exogenously added Syb2-pHluorin. The number of Syn1-positive presynaptic boutons containing GFP clusters already reached a peak at 10-30 min and did not further increase over time (Fig.3C), suggesting that Syb2-containing EVs are incorporated directly into the synapse and that there is little to no contribution of extra Syb2-pHluorin being transported from the soma in the timescale of 10 min to 2 hours.

Exogenous Syb2, added via EVs, augments inhibitory neurotransmission by increasing the propensity of synaptic vesicles to fuse (Vilcaes et al., 2021). So next we asked whether the presynaptic clusters of GFP represented endosomes or synaptic vesicles. To answer this question we evaluated the number, size and spatial distribution of GFP clusters. Most (88%) of the Syn1 presynaptic boutons only contained one GFP cluster (Fig.3D), which was found in close proximity to the plasma membrane (defined as the distance to the border of the Syn1 cluster; 52.9% at <50 nm and ∼45% at <150 nm; Fig.3E). This distribution resembles the distribution of synaptic vesicles, where the highest density was found closer to the active zone membrane (Schikorski, 2014). Average minor and major axis (diameter) of Syn1 clusters were 0.5 and 1 µm, respectively (Fig.3F), supporting that segmented clusters correspond to presynaptic boutons. Syb2-pHluorin (GFP) clusters were smaller, with mean minor and major axis of 0.2 and 0.4 µm (Fig.3F). Due to the small size and high density of synaptic vesicles, combined with the artifacts in determining size when using antibodies for detection, we cannot differentiate if larger Syb2-pHluorin (GFP) clusters represent groups of synaptic vesicles or small recycling endosomes, similar to previously described biogenic precursors (Régnier-Vigouroux et al., 1991; Watson et al., 2023). However, 6% of Syb2-pHluorin clusters had both diameters smaller than 100 nm, suggesting the detection of putative individual synaptic vesicles. Taken together, using STORM, we uncovered that Syb2-pHluorin incorporated from exogenously added EVs is localized to small vesicles in close proximity to the presynaptic plasma membrane, possibly ready to fuse in response to neuron activity.

### EVs as a new method to deliver Syb2-pHluorin to presynaptic terminals for trafficking experiments without overexpression artifacts

Our confocal and STORM experiments suggested that most of EV-delivered Syb2-pHluorin molecules resided in internal organelles, so we next corroborated this observation by measuring the ratio of pHluorin within acidic organelles compared to the neuronal plasma membrane. Surface and total fluorophore levels were measured by quenching with acidic buffer of pH=5.5 or perfusing 50 mM NH_4_Cl pH=9 solution, respectively (Fig.4A-B, also see (Sankaranarayanan et al., 2000)). The ratio of internal Syb2-pHluorin was then estimated by subtracting surface signal from total fluorescence intensity (Fig.4B). The majority (>75%) of Syb2-pHluorin incorporated into neurons from EVs resided internally, possibly in trafficking organelles (Fig.4A-B). Taken together with the STORM measurements, these results indicate that presynaptic boutons only received few copy number of Syb2-pHluorin molecules that were mostly located in one internal trafficking organelle close to the presynaptic plasma membrane. To test whether these organelles are fusion competent we stimulated neurons at low and high frequency to monitor Syb2-pHluorin activity-dependent exo-endocytosis (Fig.4A and 4C; also see (Chanaday & Kavalali, 2018, 2021)). As a negative control and threshold for detection of fusion events, we used dissociated neurons from SNAP-25 knock-out (KO) mice, which have almost complete loss of neurotransmission (Fig.4C). As expected, less than 10% of SNAP-25 KO presynaptic boutons responded to high frequency stimulation (train of 200 action potentials – APs – at 40 Hz), while 20 to 40% of wild-type (WT) synapses showed a peak in fluorescence corresponding to fusion and endocytosis of Syb2-pHluorin (Fig.4D-E). Due to the low number of synapses and synaptic vesicles receiving exogenous Syb2-pHluorin from EVs, probability of detecting fusion in response to single APs was very low (Fig.4F), but detectably larger in WT cultures compared to SNAP-25 KO (Fig.4F). In some synapses, 2 or 3 individual fusion events were detected during the recording (Fig.4F), corroborating our STORM measurements that indicated larger or multiple clusters, i.e. groups of more than one synaptic vesicle positive for Syb2-pHluorin, in some presynaptic boutons. Thus, EVs sparsely deliver Syb2 directly to presynaptic boutons leading to rapid (<30 min) integration of exogenous Syb2 into a few fusion-competent synaptic vesicles.

**Figure 4.**
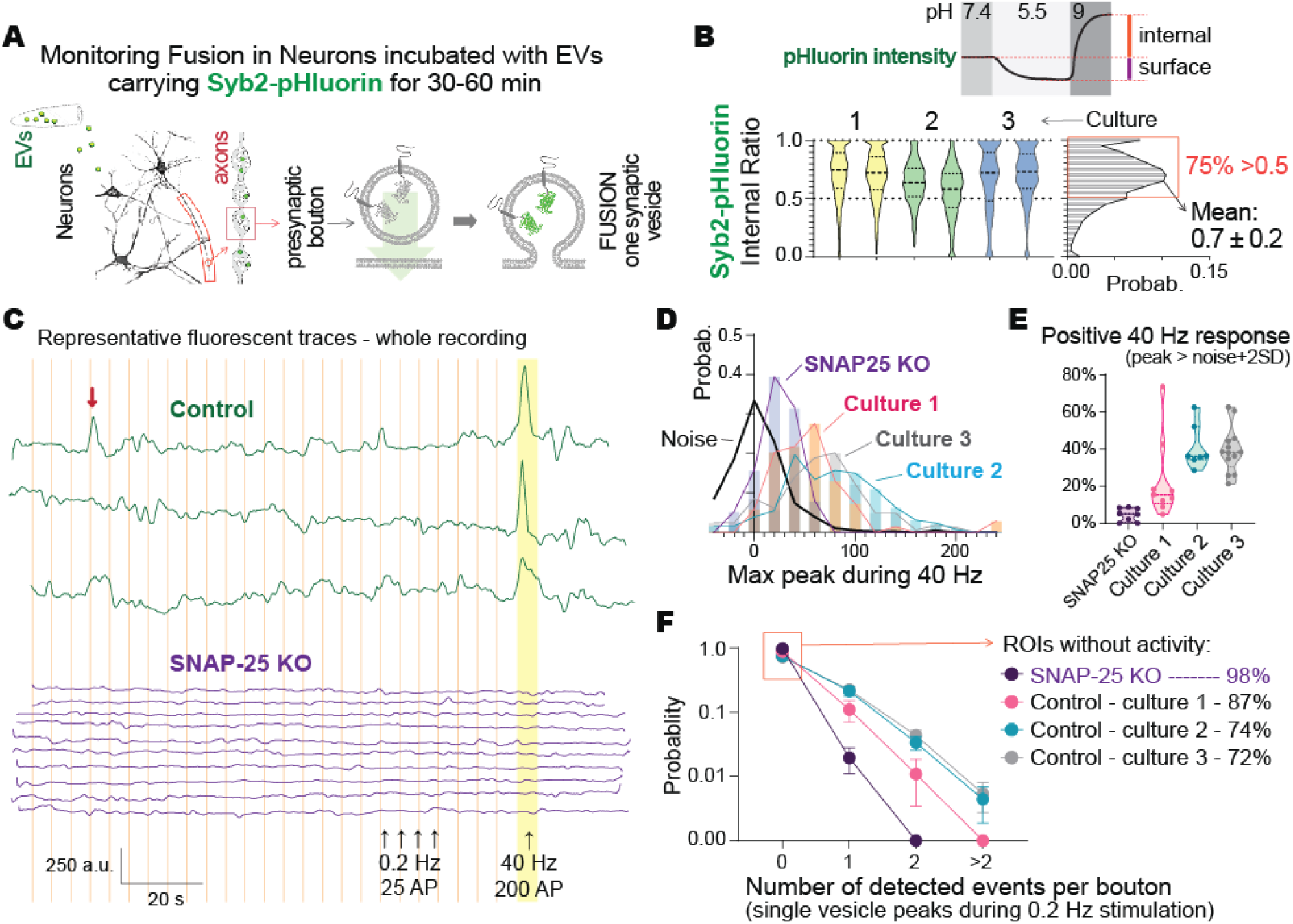
EVs sparsely deliver Syb2-pHluorin to fusion-competent presynaptic vesicles. **A**. Experimental procedure. **B**. Calculation of internal versus surface localization of Syb2-pHluorin. **C**. Fluorescence traces of control (WT) and SNAP-25 KO synapses incubated with Syb2-pHluorin EVs. **D**. Histogram of maximal fluorescence amplitude after 40 Hz stimulation. Noise + 2*SD was used as a threshold to define positive responses. **E**. Percentage of synapses that responded to 40 Hz stimulation, based on the threshold set in C. **F**. Probability of detection of single synaptic vesicle fusion events during low frequency 0.2 Hz stimulation. Each dot represents the average of all synapses from one imaging experiment, data from 3 independent cultures.

After catalyzing the release of secretory vesicles, Syb2 must be retrieved from the plasma membrane in order to facilitate successive rounds of exocytosis. Inefficient Syb2 retrieval thus leads to depression of neurotransmitter release and synaptic transmission during sustained neuron activity (Rajappa et al., 2016). Previous work tracking the fusion and endocytosis of Syb2 was performed by overexpressing Syb2-pHluorin, which leads to abnormal subcellular distribution including increased plasma membrane levels and suboptimal retrieval, primarily due to imbalance in the ratio of Syb2 to its trafficking partner Synaptophysin-1 (Gordon et al., 2016; Pennuto et al., 2003). Here we demonstrate that EVs can be used to effectively and rapidly deliver Syb2-pHluorin directly to axons, this exogenous Syb2-pHluorin is incorporated into functional secretory vesicles but only in a small subset and in very low levels, thus we next hypothesize that Syb2-pHluorin delivered via EVs can be used to track Syb2 trafficking at presynaptic terminals in the absence of overexpression artifacts. To validate that there is no overexpression of Syb2 levels, we first estimated the number of Syb2-pHluorin molecules (quanta or q) involved in each fusion event in response to single APs by fitting the distribution of amplitudes with a Gaussian mixture function (Fig.5A-C). This analysis showed that most detected vesicles carried between 1 and 4 copies of the probe (Fig.5C), which also suggests that only the smallest clusters observed via STORM are fusion competent, while the larger clusters may represent recycling or trafficking intermediaries. Considering that each synaptic vesicle carries ∼70 copies of Syb2, the exogenous addition of 1-4 copies of Syb2-pHluorin is well below endogenous levels and thus not overexpressed. We next investigated the fate of Syb2-pHluorin after fusion using our previously validated methods (Chanaday & Kavalali, 2018, 2021). We defined the kinetics of endocytosis as the time Syb2-pHluorin resides at the cell surface, exposed to extracellular pH (7.4) before being retrieved, which is calculated by measuring the duration of the dwell time in fluorescence after fusion (Fig.5A). After endocytosis, the endosome or vesicle is acidified (pH∼5), quenching the fluorophore and causing a decay in fluorescence. By analyzing fusion events from individual vesicles, we can thus deconvolve endocytosis – dwell time – from acidification – decay time. Calculated dwell times pointed to very fast retrieval of Syb2-pHluorin after fusion (Fig.5D-E). Fitting of logarithmic histograms with Gaussian mixture models predicted the presence of two populations with characteristic endocytic times of 700 ms and 1.5 s, corresponding to our previously reported ultrafast and fast endocytic pathways (Chanaday & Kavalali, 2018, 2021). However, we did not detect slow endocytosis (>5s), suggesting this previously described mode of synaptic vesicle retrieval is either a very low probability event or a consequence of overexpression. Newly exocytosed Syb2 was previously proposed to undergo confined diffusion out of the presynaptic active zone (Gimber et al., 2015), with fusion being compensated by retrieval of preassembled clusters of synaptic vesicle and endocytic proteins at the periphery of the active zone (Del Signore et al., 2023; Hua et al., 2011; Imoto et al., 2022). Since Syb2-pHluorin delivered via EVs resided mainly in internal organelles (Fig.4B), our system presented the ideal opportunity to track the destination of newly fused Syb2, i.e. whether it stays at the surface or it is rapidly retrieved, in the absence of confounding effects from surface signal. We defined the efficiency of retrieval as the ratio or fraction of the amplitude of the decay phase (peak back to baseline) – which depends on the number of Syb2-pHluorin molecules that were retrieved, respect to the amplitude of the fusion event (baseline to peak) – which corresponds to the number of fused Syb2-pHluorin (Fig.5F). The majority (>75%) of the fusion events showed quantal retrieval, i.e. a ratio of 1.0±0.2 (Fig.5F), suggesting that for most fusion events all the fused probes were subsequently endocytosed. The remaining fusion events showed partial (14%) or no retrieval (8%; Fig.5F), indicating that in some cases the fused molecules are not instantly retrieved and remain at the plasma membrane, and a different portion of the membrane may undergo compensatory endocytosis. Taken together, by using EVs to sparsely deliver Syb2-pHluorin to synaptic vesicles we show that the main mode of Syb2 retrieval during low frequency activity consists of ultrafast and highly efficient endocytosis of newly fused Syb2.

**Figure 5.**
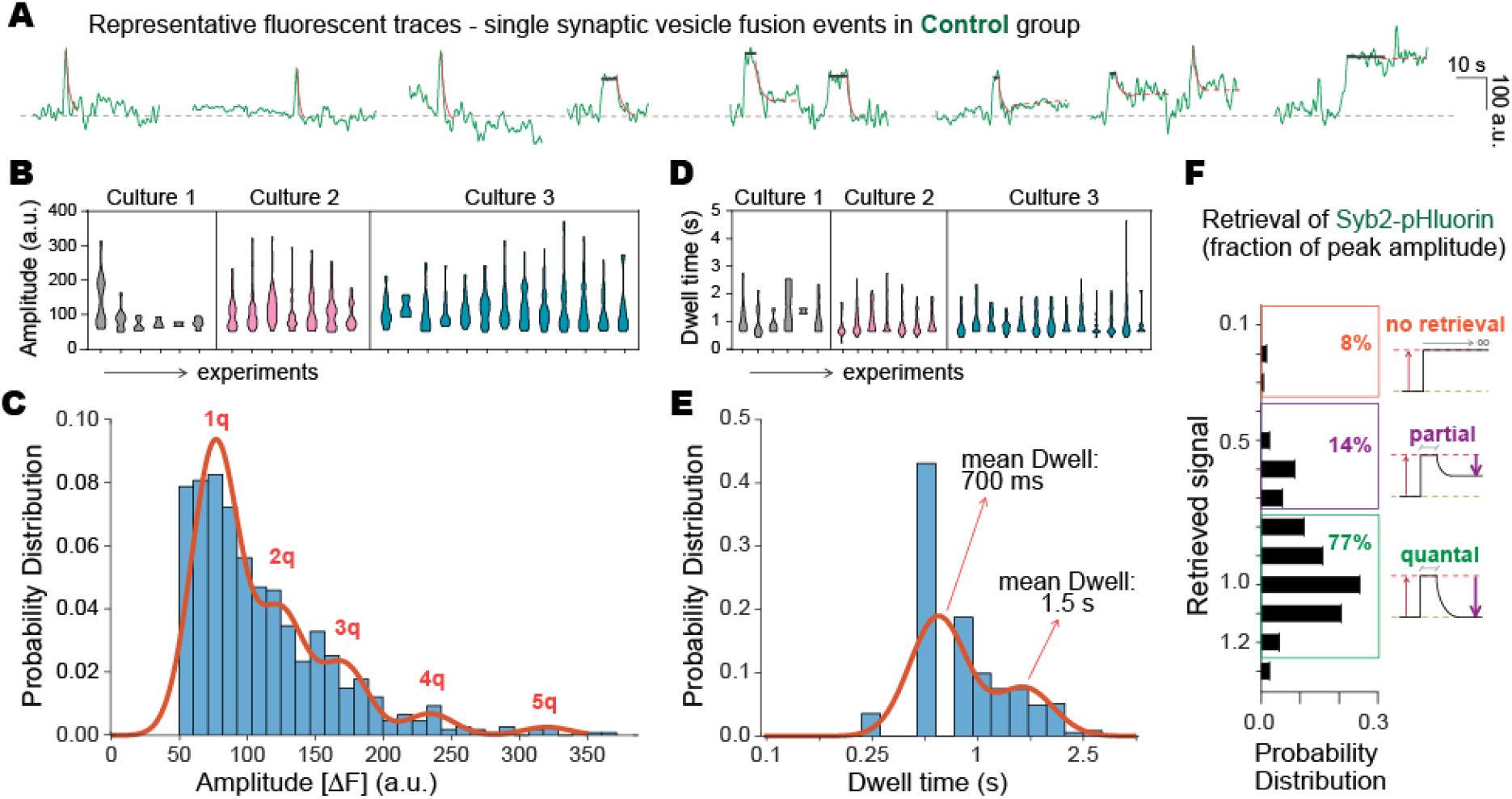
Newly fused Syb2 is rapidly and efficiently retrieved during synaptic activity. **A**. Fluorescence traces of single vesicle fusion events. **H-I**. Amplitudes and **J-K**. Dwell times of single synaptic vesicle fusion events. For G and I, distributions were fitted with a Gaussian mixture model. **L**. Retrieval of Syb2-pHluorin, calculated as the ratio of total fluorescence decay after fusion over the event’s amplitude.

## DISCUSSION

In the current study we took advantage of our previous discovery of Syb2 being secreted by neurons via EVs to answer two central questions in Syb2 trafficking: the uptake mechanism and location of Syb2-containing EVs by neurons; and the fate of newly fused Syb2 molecules during low frequency synaptic activity. We demonstrate that Syb2-containing EVs are incorporated throughout the neuron, including soma and axon, via a Clathrin-independent and Dynamin-independent pathway. We next show that EVs can be used to sparsely deliver Syb2-pHluorin to synaptic vesicles and track their activity-driven fusion and endocytosis without overexpression artifacts. Using this approach we show that following the fusion of single synaptic vesicles the main mode of Syb2 retrieval is ultrafast and highly efficient, supporting a model where synaptic vesicle molecules remain clustered and are rapidly cleared out of the plasma membrane.

### Uptake of Syb2-containing EVs into Neurons

Mounting evidence suggests that EVs are incorporated into various target cells primarily via endocytosis, including Clathrin-mediated, Caveolin-dependent, macropinocytosis and phagocytosis (Escrevente et al., 2011; Feng et al., 2010; Fitzner et al., 2011; Mulcahy et al., 2014; Solana-Balaguer et al., 2023; Tian et al., 2010). Here we show that a specific subpopulation of neuronal EVs that carry the presynaptic protein Syb2 is internalized into hippocampal neurons via a separate route compared to the Caveolin cargo LacCer and the Clathrin cargo TF receptor. Moreover, the Dynamin inhibitor Dyngo-4a did not impede the uptake of Syb2 from EVs, demonstrating that neuronal Syb2-containing EVs internalization is Dynamin-independent. It is possible that other types of neuronal EVs that lack Syb2 follow alternative uptake and transport routes as suggested previously in other cell types (Escrevente et al., 2011; Ginini et al., 2022; Mulcahy et al., 2014; Pedrioli & Paganetti, 2020). In this context, our results investigating a specific subpopulation of neuronal EVs that carry Syb2 support the premise that different subtypes of EVs, depending on their molecular composition, are internalized via separate routes. We additionally show that EVs uptake is very rapid, peaking at 10-30 min, and occurs throughout neuron compartments, including the soma and axon. Previous work in other neuron types also highlighted that EVs uptake seem to mainly occur in the neuron soma (Forero et al., 2024), which may facilitate the delivery of genetic materials and transcription factors to the nucleus. Our data shows that Syb2-containing EVs are also directly incorporated by axons and integrated into functional synaptic vesicles thus influencing synaptic transmission, further supporting the premise that the subcellular location of EV internalization is crucial for fast delivery of its cargoes and for determining their functional effect. After internalization, Syb2 delivered via EVs follows an intra-neuronal transport route that converges with that of LacCer and is maintained completely segregated from TF receptor trafficking, showing that transport of EV components is not random but follows specific pathways possibly mimicking the normal transport mechanisms of the endogenous molecules.

Our experiments show that Syb2-containing EVs are internalized independently of Dynamin. A number of Dynamin-independent endocytic pathways were proposed in non-neuronal cell types for EVs uptake, including macropinocytosis and lipid raft mediated (Escrevente et al., 2011; Ginini et al., 2022; Mulcahy et al., 2014; Pedrioli & Paganetti, 2020). While there is no evidence of these forms of endocytosis specifically mediating EVs uptake in neurons, macropinocytosis does occur in neurons and it mediates processes including axon growth and integration of large protein aggregates during neurodegeneration (Powers et al., 2022; Zeineddine & Yerbury, 2015). Lipid raft mediated endocytosis is of particular interest since neurons and EVs are enriched in glycolipids (gangliosides) associated with these less-fluid membrane domains (Monyror et al., 2025; Yuyama et al., 2014). Another interesting candidate for Dynamin-independent EVs uptake is CLIC/GEEC endocytosis, since it mediates endocytosis of large portions of plasma membrane in various polarized cell types selectively at regions rich for gangliosides and cholesterol (Shafaq-Zadah et al., 2020). Discerning among these uptake routes will be experimentally challenging, however, since they have overlapping molecular machinery – among them and with other transport routes – and there is no specific pharmacology.

### Using EVs for sparse labeling of synaptic vesicles with Syb2-pHluorin

Previous studies proposed several endocytic mechanisms to retrieve synaptic vesicles during neuron activity. One possibility is diffusion of synaptic vesicle molecules after full collapse fusion, and compensatory ultrafast or bulk endocytosis of a different portion of membrane in the periactive zone (Chanaday et al., 2019; Del Signore et al., 2023; Gimber et al., 2015; Li & Murthy, 2001; Ogunmowo et al., 2023; Wu et al., 2014). Another option is that synaptic vesicle molecules remain clustered after fusion and are rapidly recaptured and retrieved (Alabi & Tsien, 2013), a mechanism recently called “kiss-shrink-run” (Tao et al., 2025). Or a combination of both scenarios. Previous discrepancies arise from using different methodologies, with variable spatial and temporal resolutions. Another caveat is the use of overexpression of pHluorin-tagged proteins, which causes mislocalization, aberrant lateral diffusion and defects in retrieval from the plasma membrane (Gordon et al., 2016; Pennuto et al., 2003). Here, we overcame these caveats by using EVs to deliver Syb2-pHluorin to synapses. Our results show that fast and ultrafast endocytosis mediate most of the retrieval of synaptic vesicles during low-frequency firing. The majority of these events retrieve all the fused molecules, indicating that synaptic vesicle components may remain clustered for fast recapture and retrieval. The remaining events show partial to no retrieval, suggesting the coexistence with diffusion and compensatory endocytosis, and agreeing with recent measurements using orthogonal methods (Tao et al., 2025). These results highlight the richness and complexity of mechanisms regulating synaptic vesicle recycling at synapses. This recycling diversity may have evolved to provide resilience such that synapses can continue to function even if endocytic mechanisms are compromised. This work opens the door for future applications of EVs for selective delivery of molecules for the study of transport and signaling in neurons.

A number of molecular mechanisms have been found to support synaptic vesicle recycling, including Clathrin-mediated endocytosis and regeneration from endosomes, Dynamin-dependent fast and ultrafast retrieval mediated by Formins and/or Synaptojanin and Endophilin, and Dynamin-independent endocytosis (Afuwape et al., 2024; Chanaday et al., 2019; Gan & Watanabe, 2018; Ivanova & Cousin, 2022; Shin et al., 2021; Soykan et al., 2017). In the future it would be interesting to elucidate whether the Dynamin-independent integration of Syb2-contanining EVs we demonstrate here is hijacking the same Dynamin-independent mechanism that endogenous synaptic vesicles utilize for their fast recycling at presynaptic terminals during neuron activity (Afuwape et al., 2024).

## ABBREVIATIONS

Syb2: Synaptobrevin-2
EVs: extracellular vesicles
MAP2: microtubule-associated protein 2
TF: Transferrin-Alexa Fluor 647
LacCer: BODIPY-lactosylceramide
Syn1: Synapsin-1

## Author Contributions

V.H., A.A.V. and N.L.C. conceived the study. V.H., A.A.V., Q.T., S.H. and N.L.C. designed, performed, analyzed and interpreted the experiments. V.H., A.A.V. and N.L.C. made the figures and wrote the initial draft of the manuscript. V.H., A.A.V., Q.T., S.H., M.L. and N.L.C. wrote the final version of the manuscript. M.L. and N.L.C. secured the funding for this research.

## Acknowledgements

This work was supported by the University of Pennsylvania and the Margaret Q. Landenberger Research Foundation Award to N.L.C. We thank the CDB Microscopy Core (RRID SCR_022373) and the Extracellular Vesicle Core Facility (Penn Vet.; RRID:SCR_022444) of the University of Pennsylvania for the use of their instruments.

